# The mechanism of full activation of tumor suppressor PTEN at the phosphoinositide-enriched membrane

**DOI:** 10.1101/2021.01.06.425565

**Authors:** Hyunbum Jang, Iris Nira Smith, Charis Eng, Ruth Nussinov

**Affiliations:** Computational Structural Biology Section, Frederick National Laboratory for Cancer Research in the Laboratory of Cancer Immunometabolism, National Cancer Institute, Frederick, MD 21702, U.S.A.; Genomic Medicine Institute, Cleveland Clinic Lerner Research Institute, Cleveland, OH 44195, U.S.A.; Center for Personalized Genetic Healthcare, Cleveland Clinic Community Care and Population Health, Cleveland, OH 44195, U.S.A.; Department of Genetics and Genome Sciences, Case Western Reserve University School of Medicine, Cleveland, OH 44106, U.S.A.; Germline High Risk Cancer Focus Group, Comprehensive Cancer Center, Case Western Reserve University School of Medicine, Cleveland, OH 44106, U.S.A.; Department of Human Molecular Genetics and Biochemistry, Sackler School of Medicine, Tel Aviv University, Tel Aviv 69978, Israel

**Keywords:** phosphatase, dephosphorylation, signaling lipid, phosphoinositide, PIP_3_, PIP_2_, protein-membrane interaction, Ras

## Abstract

Tumor suppressor PTEN, the second most highly mutated protein in cancer, dephosphorylates signaling lipid PIP_3_ produced by PI3Ks. Excess PIP_3_ promotes cell proliferation. The mechanism at the membrane of this pivotal phosphatase is unknown hindering drug discovery. Exploiting explicit solvent simulations, we tracked full-length PTEN trafficking from the cytosol to the membrane. We observed its interaction with membranes composed of zwitterionic phosphatidylcholine, anionic phosphatidylserine, and phosphoinositides, including signaling lipids PIP_2_ and PIP_3_. We tracked it’s moving away from the zwitterionic and getting absorbed onto anionic membrane that harbors PIP_3_. We followed it localizing on microdomains enriched in signaling lipids, as PI3K does, and observed PIP_3_ allosterically unfolding the N-terminal PIP_2_ binding domain, positioning it favorably for the polybasic motif interaction with PIP_2_. Finally, we determined PTEN catalytic action at the membrane, all in line with experimental observations, deciphering the mechanisms of how PTEN anchors to the membrane and restrains cancer.

## Introduction

Tumor suppressor phosphatase and tensin homolog (PTEN) dephosphorylates phosphatidylinositol 3,4,5-trisphosphate (PIP_3_) to phosphatidylinositol 4,5-bisphosphate (PIP_2_) (Chalhoub and Baker, 2009). Signaling lipid PIP_3_, a major product of phosphatidylinositol 3-kinases (PI3Ks), recruits a subset of proteins, phosphoinositide-dependent protein kinase 1 (PDK1) and AKT to the plasma membrane, leading to cell growth, survival, and migration (Fruman et al., 2017). Both PTEN and PI3K are among the most frequently mutated proteins in malignancies, acting as negative and positive regulators in the PI3K/AKT/mTOR pathway (Chalhoub and Baker, 2009; Song et al., 2012). Countering PI3K lipid kinase activity, PTEN lipid phosphatase suppresses the level of PIP_3_ in the cell membrane. In many cancers, elevation of PIP_3_ level is linked to increased activity of PI3K in the oncogenic state (Fruman et al., 2017). In human malignancies, both sporadic (Yehia et al., 2019) and heritable (Yehia et al., 2018) loss of PTEN function by respectively, somatic and germline mutations, deletions, posttranslational modifications, and phosphorylation lead to PIP_3_ accumulation in the cell membrane, boosting unrestrained PI3K-stimulated cell proliferation (Alvarez-Garcia et al., 2019; Dillon and Miller, 2014; Kotelevets et al., 2020; Song et al., 2012).

With 403 amino acids, PTEN is composed of the N-terminal PIP_2_ binding domain (PBD aka PBM, PIP_2_ binding motif, residues 1-15), the phosphatase domain (residues 16-185), and the C2 domain (190-350), followed by the carboxy-terminal tail (CTT) including the PDZ binding motif (PDZ-BM, 401TKV403) at the C-terminal end (Figure 1). The phosphatase domain is key in PTEN’s catalytic activity with the P loop (residues 123-130) containing the catalytic signature motif, HCxxGxxR (where x is any amino acid), commonly found in the active sites of other protein tyrosine phosphatases (Denu et al., 1996; Lee et al., 1999). Within the motif, Cys124 is crucial for transferring a phosphate group from the inositol of PIP_3_ during the catalytic activity (Xiao et al., 2007). Reversible oxidation involving disulfide formation between Cys71 and Cys124 at the active site inhibits the phosphatase activity of PTEN (Lee et al., 2015). In the cytoplasm, PTEN exists as a monomer and can even form a homodimer *in vitro* (Heinrich et al., 2015). The disordered CTT is key to the regulation of the monomeric PTEN state in the cytosol through the phosphorylation of the serine-threonine cluster (Ser380, Thr382, Thr383, and Ser385) located in it (Bolduc et al., 2013; Masson et al., 2016; Masson and Williams, 2020; Mingo et al., 2019; Vazquez et al., 2000). Cytosolic PTEN monomers exhibit a “closed-closed” conformation with phosphorylated CTT (“closed” CTT covers the membrane binding interface of the phosphatase and C2 domains, and “closed” PBD nestles against the phosphatase domain), and an “open-closed” conformation with dephosphorylated CTT (Malaney et al., 2013; Rahdar et al., 2009; Ross and Gericke, 2009) (“open” CTT exposes the membrane binding interface of the phosphatase and C2 domains). The dephosphorylated CTT releases the autoinhibition in the active site, facilitating the translocation of PTEN to the membrane (Nussinov et al., 2020). At the membrane, PTEN yields an “open-open” conformation, finally releasing its PBD. Experimental data indicated that electrostatic interactions between the positively charged residues in the PBD and the C2, and the negatively charged anionic lipids play a crucial role in the recruitment of PTEN to the membrane (Denning et al., 2007; Nguyen et al., 2014b; Walker et al., 2004). In particular, the short N-terminal PBD is important for controlling PTEN membrane localization. This established description has provided a useful phenomenological overview of PTEN. However, the mechanistic details of the membrane association, membrane-bound conformations, and catalytic conformational changes of PTEN at atomic resolution, critical for gaining functional insight into the workings of this tumor suppressor and possible new venues for drug discovery, are still unknown.

**Figure 1.**
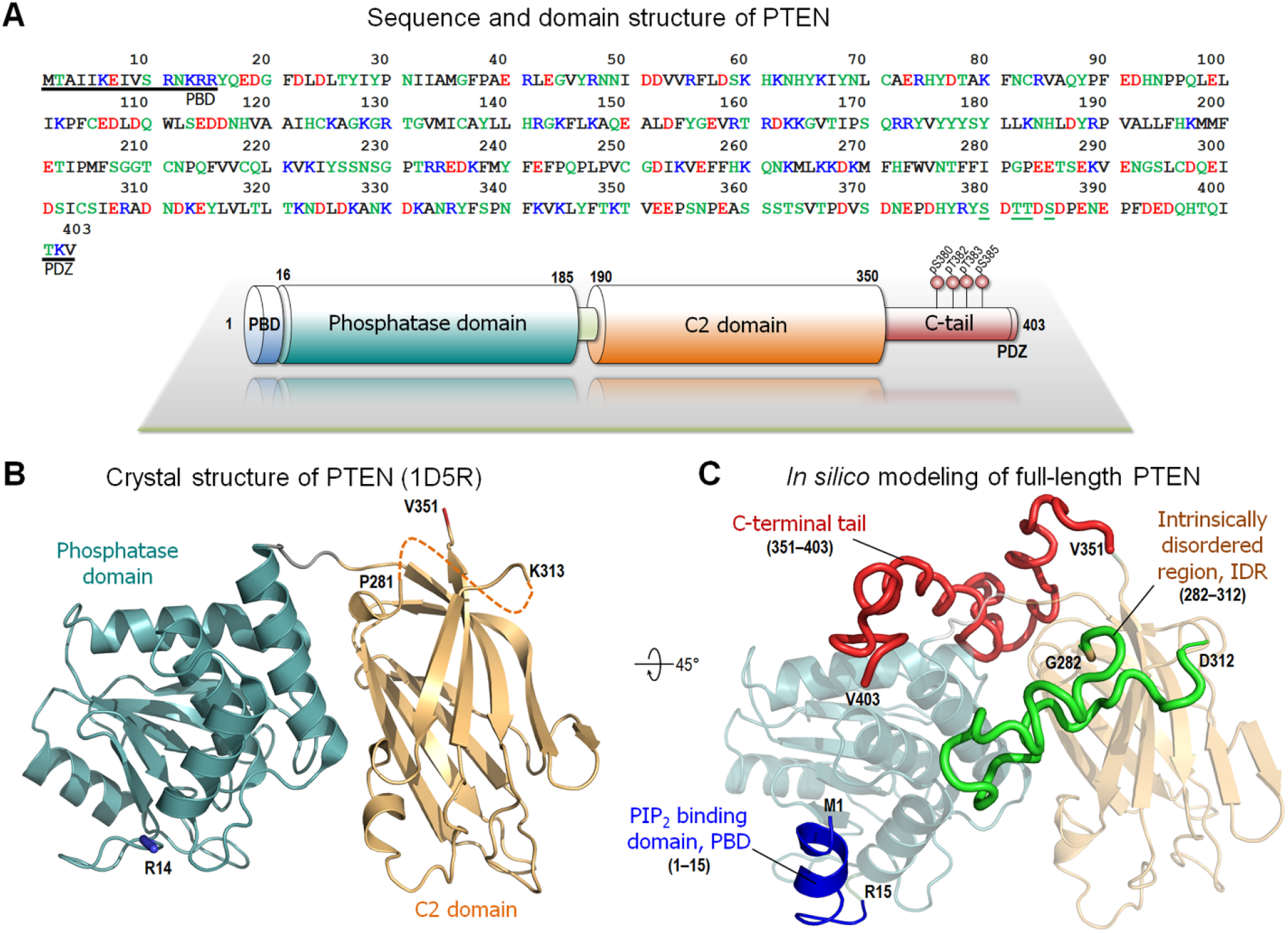
PTEN sequence and structure. (A) The sequence and domain structure of PTEN. In the sequence, the underlined residues highlight the PIP_2_ binding domain (PBD) at the N-terminal and the PDZ binding motif at the C-terminal regions. In the domain structure, the phosphorylated serine-threonine cluster (residues 380-385) in the C-terminal tail is marked. (B) The crystal structures of PTEN (PDB code: 1D5R). The marked residues denote the missing regions. (C) *In silico* model of the full-length PTEN structure. The missing regions in PTEN, the PBD (residues 1-15), the interdomain region (IDR, residues 282-312), and the C-terminal tail (residues 351-403) were constructed by the I-TASSER program (Roy et al., 2010; Yang et al., 2015; Yang and Zhang, 2015).

Here, we aim to provide such mechanistic understanding of PTEN at the membrane. Our study is extensive. We performed explicit solvent MD simulations on full-length, wild-type PTEN interacting with lipid bilayers. The bilayer systems mainly consist of zwitterionic, 1,2-dioleoyl-sn-glycero-3-phosphocholine (DOPC) and anionic, 1,2-dioleoyl-sn-glycero-3-phosphoserine (DOPS) lipids. In addition, the signaling phosphoinositides, PIP_2_, and PIP_3_, were added as described in Table S1. The initial conformation of PTEN at the membrane is the “open-closed” conformation with dephosphorylated CTT. Dephosphorylation of CTT uncovers the active site in the phosphatase domain, permitting PTEN to exert its catalytic function targeting PIP_3_ in the membrane (Nussinov et al., 2020). In our studies, PTEN stably anchored in the anionic membrane with both signaling lipids, and with the two positively charged loops, one from the phosphatase domain and the other from the C2 domain, inserted into the amphipathic interface of the lipid bilayer. Remarkably, PTEN evolved into the “open-open” conformation, spontaneously releasing the PBD from the phosphatase domain to nearby PIP_2_ in the membrane. The additional anchor by PBD further stabilized the PTEN membrane association. Effective membrane localization is crucial for PTEN to exert its lipid phosphatase activity, which leads to the inhibitory effect on signaling via the PI3K/AKT/mTOR pathway. Our studies identify the detailed mechanisms deciphering atomistic interactions of how PTEN anchors to the membrane and elucidate the vital mechanistic role of PTEN in restraining cancer.

## Results

### PBD targets PIP_2_ via conformational change and membrane localization

MD simulations were performed on full-length PTEN interacting with an anionic lipid bilayer composed of DOPC, DOPS, PIP_2_, and PIP_3_ lipids (hereafter denoted as PP system, Table S1). The anionic bilayer system was constructed through random distribution of the anionic lipids on the plane of the bilayer, except PIP_3_ which was placed nearer the protein. PTEN was initially placed on the lipid bilayer with its putative membrane-binding interface facing toward the bilayer surface as suggested by the crystal structure (Lee et al., 1999). An asymmetric profile of electron density across the bilayers indicates that the placement of the protein depletes the electron density due to the volume occupied by the bulky waters (Figure S1A). In the profile, the distance between two peaks determines the bilayer thickness. The initial configuration of the PTEN-bilayer system ensures that the phosphatase and C2 domains are placed on the top of the bilayer surface without inserting the protein backbone into the bilayer. In this initial configuration, both the PBD and the C-tail do not directly interact with the lipid bilayer (Figure S1B). However, during the simulation, we observed that PTEN gradually diffuses into the bilayer and fully relaxed in the lipid environment (Figure 2A). The full relaxation process for the PTEN-bilayer system requires several hundred nanoseconds, which is comparatively longer than that for the protein-bulky water system. For most of the anionic bilayer systems (PS, P2, P3, and PP systems, defined in Table S1), we observed that the measured quantities such as the deviations of PTEN domains from the membrane surface generally reached constant values at ∼400 ns. For PTEN in the PP system, the phosphatase domain retains its structural integrity with averaged root-mean-squared-deviation (RMSD) of 1.7 ± 0.2 Å with respect to the starting point. While the C2 domain preserves a β-sandwich structure, its RMSD increases to 4.4 ± 0.9 Å due to fluctuations in the unstructured IDR (intrinsically disordered region; residues 282-312). Large fluctuations are prominent in the unstructured regions including the C-terminal region as observed in the residual RMS-fluctuations (Figure 2B). Of interest is the large conformational change of PBD. The initial α-helical motif in PBD is rapidly disrupted and converted into an unstructured chain (Figure S1C). Notably, after ∼400 ns, the unstructured PBD spontaneously docks onto the bilayer surface establishing a secure anchor point for the phosphatase domain in the membrane (Figure 2C). The membrane anchorage of PBD is sustained by strong salt bridge interactions between the anionic lipids and the polybasic patch (Arg11, Lys13, Arg14, and Arg15), resulting in the coordination of two PIP_2_ lipids (Figure 2D). The N-terminus residue Met1 diffuses into the hydrophobic core of the lipid bilayer, inducing nearby Lys6 to localize at the amphipathic interface of the bilayer and to interact with a phosphate group of DOPC. Although our RMS-fluctuations data show that PBD has large fluctuations due to the conformational change and the membrane translocation, the dipping of the N-terminus in the bilayer reduces these fluctuations, thus enhancing the stability of the PTEN membrane interaction following completion of the PBD anchorage.

**Figure 2.**
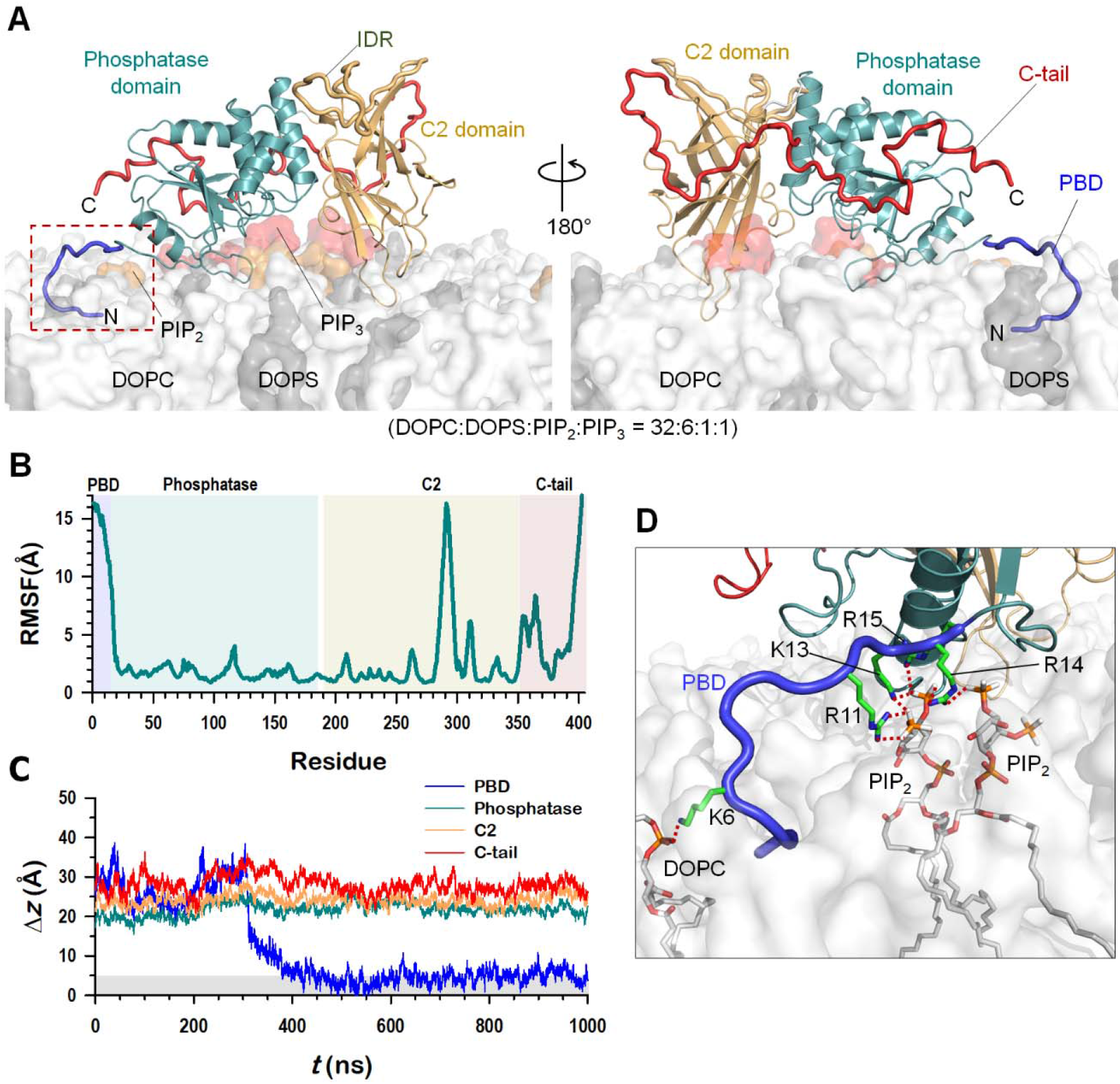
Membrane interaction of PTEN. (A) Snapshot representing the final PTEN conformation in the lipid bilayer composed of DOPC:DOPS:PIP_2_:PIP_3_ = 32:6:1:1, in molar ratio. (B) The root-mean-squared-fluctuations (RMSFs) of PTEN residues. (C) Time series of the deviations of the center of mass of individual PTEN domains from the bilayer surface. (D) Highlight showing the interaction of unfolded PBD with the lipids. Key basic residues, Lys6, Arg11, Lys13, Arg14, and Arg15 are marked.

### PTEN localization is highly sensitive to lipid composition

PTEN targets the substrate lipid PIP_3_ and co-localizes with it. The substrate is in the inner leaflet of PIP_3_-rich microdomain within the plasma membrane. Thus, precise PTEN membrane localization on the substrate is crucial for exerting its lipid phosphatase activity. To decipher how PTEN is effectively absorbed in the membrane in the presence/absence of the PIP_3_ substrate, simulations were performed for PTEN with various lipid compositions in the bilayers as outlined in Table S1. To derive results from the lipid effects, the same PTEN as in the PP system (Figure S1B) was applied for all bilayer systems, ensuring that all starting points have the same initial membrane orientation and location for the protein. To identify membrane binding sites, the probability distribution functions of membrane contacts were measured as a function of the protein residues (Figure 3). Remarkably, for PTEN in the zwitterionic DOPC bilayer (PC system), the phosphatase domain loses contacts with the membrane (Figure S2A) as indicated by the low probability of membrane contacts for both PBD and phosphatase domain. The phosphatase domain is highly elevated from the bilayer surface (Figure S2B), while the C2 domain is still attached to the bilayer, which prevents entire protein domains from flipping on the bilayer surface. However, in the presence of anionic DOPS lipid (PS system), PTEN regains its membrane attachment ability. Six loops located in the phosphatase and C2 domains are responsible for the membrane association. These are PBD-pβ1 (19DGFDL23), pβ2-pα1 (aka Arginine loop, 41RLEGVYR47), and pα5-pα6 (aka TI loop, 161RDKKGV166) loops in the phosphatase domain, and cβ1-cβ2 (205MFSGGTC211), CBR3 (260KQNKMLKKDK269), and Cα2 (aka Motif 4, 327KANKDKANR335) loops in the C2 domain. Among these, the pβ2-pα1, CBR3, and Cα2 loops were identified as membrane-binding regulatory interfaces in PTEN (Nguyen et al., 2014a; Nguyen et al., 2014b). In particular, the Lys-rich CBR3 loop is known as the phosphatidylserine binding motif for the C2 domain (Huang et al., 2012; Lee et al., 1999). In the P2 system with the addition of PIP_2_, PTEN still conserves all six loop regions for the membrane contact. However, in the P3 system with the addition of PIP_3_, the TI loop completely loses the membrane contact. The same behavior is also observed for the TI loop in the PP system with the addition of both signaling lipids, PIP_2_ and PIP_3_ (Figure 3, bottom panel). Moreover, the Cα2 loop weakens the membrane contacts. For the TI loop, the loss of the membrane contacts is closely related to the presence of PIP_3_, while it is related to the presence of PIP_2_ for the Cα2 loop. These signaling lipids move into the region underneath the interface between the phosphatase and C2 domains, bolstering PTEN on the membrane. Especially, the interaction of PIP_3_ with the phosphatase domain yields a distinct membrane orientation for the domain. We observed that PIP_3_-containing P3 and PP systems generate a similar profile of the distributions of the helix tilt angles for the helices in the phosphatase domain (Figure S3). Unlike the phosphatase domain, the C2 domain retains its membrane binding sites for all anionic bilayer systems, indicating its intrinsic feature that targets cell membrane.

**Figure 3.**
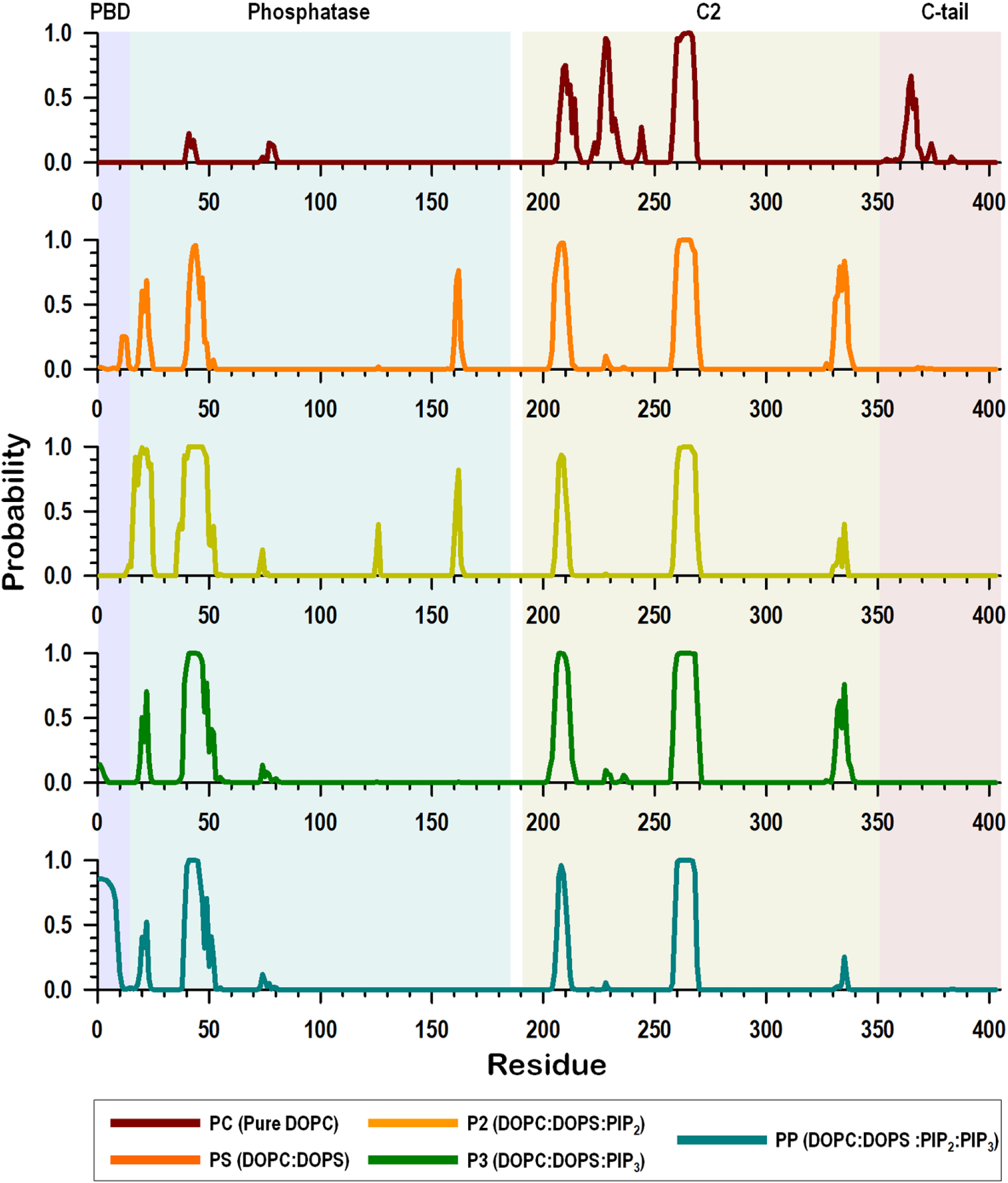
Lipid contact probability. The probability of lipid contacts for PTEN residues for different bilayer systems, including PC (pure DOPC), PS (DOPC:DOPS = 4:1, molar ratio), P2 (DOPC:DOPS:PIP_2_ = 16:3:1), P3 (DOPC:DOPS:PIP_3_ = 16:3:1) and PP (DOPC:DOPS:PIP_2_:PIP_3_ = 32:6:1:1) systems.

In the absence of the interaction between PBD and PIP_2_, the spontaneous unfolding of PBD can be observed. While the PBD retains the initial α-helical motif in the PC and PS systems, it is partially unfolded in the P2 system (Figure S2A) and completely unfolded in the P3 system (Figure S4A) in the absence of PBD–PIP_2_ interaction. The unfolding of PBD is also observed in the PP system (Figure S1C) in the presence of the PBD–PIP_2_ interaction, suggesting that PBD has an intrinsic propensity to unfold regardless of the interaction with PIP_2_. Only P3 and PP systems contain PIP_3_, suggesting that PIP_3_ at the active site may allosterically promote PBD unfolding. In the P3 system, the unfolded PBD emerges at ∼350 ns, which is relatively longer than the PP system with ∼40 ns. We observed abrupt changes in the deviation of the center of mass of PBD from the bilayer surface, indicating that the N-terminus flips the location back and forth with respect to the phosphatase domain (Figure S4B). In the first flip event, the PBD failed to secure the interaction with the membrane due to unavailability of anionic lipids at the flipping site. However, in the presence of anionic lipids near the PBD binding site, the polybasic motif 11RNKRR15 is able to interact with DOPS as in the PS system (Figure S4C) and PIP_2_ as in the P2 system (Figure S4D) even though both systems have the folded α-helical PBD sequestered by the phosphatase domain. We expect that the folded PBD eventually gets unfolded due to its intrinsic propensity to be an unstructured domain (Redfern et al., 2008). Among the polybasic residues in PBD, Lys13 and Arg14 appeared to be key residues interacting with anionic lipids. It was reported that K13E mutation impairs the ability of PTEN to bind anionic lipid bilayer (Walker et al., 2004).

### Strong salt bridge interactions between key basic residues and phosphoinositide lipids conduce PTEN membrane localization

To corroborate the effects of phosphoinositide lipids on the membrane localization of PTEN, we trace the interactions of basic residues in both phosphatase and C2 domains. PTEN contains a high population of basic residues (Arg and Lys) at the membrane binding interface, 29 out of the total 54 basic residues participate in interactions with lipids. These basic residues are mainly located at the PBD and the loop regions in the phosphatase and C2 domains, forming salt bridges with the phosphate group of the lipids and the head groups of the anionic lipids. Both the PBD and phosphatase domain are Arg-rich, while the C2 domain is Lys-rich (Figure 4A). The high probability of salt bridge formation indicates that Arg favors PIP_2_, while Lys favors PIP_3_ (Figure 4B). Those Arg residues, Arg14 (in PBD), Arg161 (in phosphatase), and Arg335 (in C2) exhibited high probability of forming salt bridges with PIP_2_ (Figure S5A). In contrast, those Lys residues, Lys125, Lys128, and Lys164 in the phosphatase domain, and Lys221, Lys223, Lys267, Lys269, and Lys330 in the C2 domain exhibited high probability of forming salt bridges with PIP_3_. While PIP_2_-favored Arg residues are located proximal to the bilayer, PIP_3_-favored Lys residues including a key Arg residue, Arg130, are located distal to the bilayer surface. Basic residues with high probability of forming salt bridges with PIP_3_ have large deviations from the bilayer surface (Figure 4C), suggesting that PIP_3_ favors distal basic residues in the salt bridge interactions. Such basic residues, Lys125, Lys128, and Arg130 can be found in the P loop, which is located at ∼15 Å, largely elevated from the bilayer surface. In salt bridge interactions with DOPC and DOPS, basic residues exhibit a similar pattern (Figure S5B), indicating that salt bridges with high probability involve basic residues in loop regions that inserted into the amphipathic bilayer interface. These are the pβ2-pα1 and CBR3 loops in the phosphatase and C2 domains, respectively. In this case, the basic residues can be located close to the bilayer surface and form salt bridges with the phosphate and acidic head groups of bulky lipids. Arg41 in the pβ2-pα1 loop strongly interacts with the lipids as observed in experiments (Nguyen et al., 2014b). The CBR3 loop in the C2 domain is embedded in the amphipathic interface, promoting its basic residues to form salt bridges with DOPS with relatively high probability as compared to other anionic lipids. Each anionic lipid has binding preference for PTEN binding sites without overlapping each other (Figure 4B and Figure S5), suggesting that PTEN synergistically interacts with the lipids as observed in the experiments (Campbell et al., 2003; Redfern et al., 2008).

**Figure 4.**
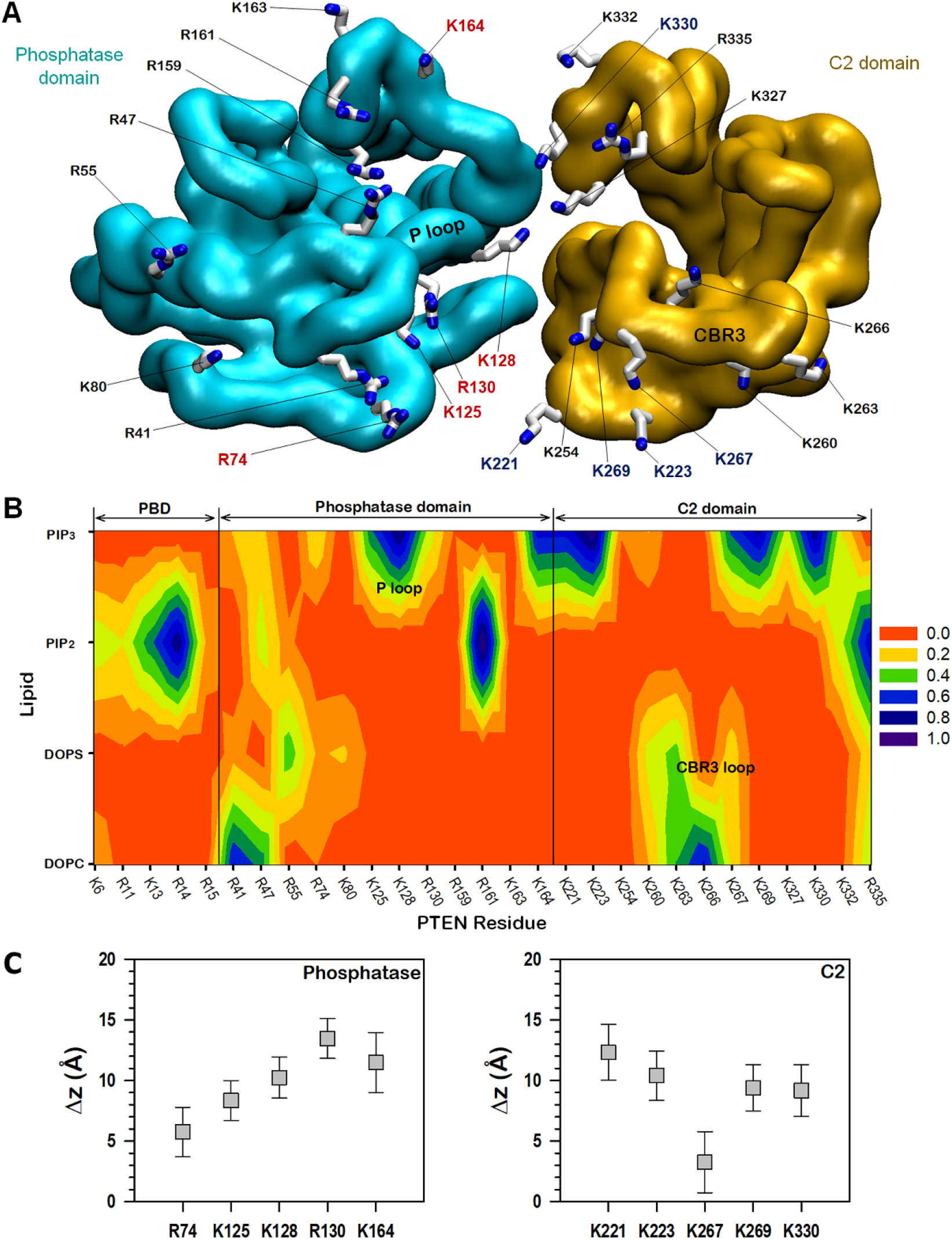
Salt bridge interactions of PTEN. (A) Mapping of the basic residues on the membrane-binding surface of the phosphatase and C2 domains. PIP_3_-favored residues in the phosphatase domain are colored red, and those in the C2 domain are colored blue. (B) Contour map of the probabilities of salt bridge formations between the key basic residues at the membrane-binding interface and the lipids. Salt bridge is calculated between the nitrogen atoms in the sidechains of basic residues and the oxygen atoms in the phosphate group of all lipids and the inositol head group of phosphoinositides with the cutoff distance of 3.2 Å. (C) Averaged deviations of the amide nitrogen in the sidechains of Arg and Lys residues from the bilayer surface for the PIP_3_-favored residues in the phosphatase (left panel) and C2 (right panel) domains.

### Substrate coordination and conformational change of the P loop implicate PTEN catalytic activity

In the lipid phosphatase activity of PTEN, the P loop at the active site plays a significant role in transferring a phosphate group from the PIP_3_ substrate. To recruit the substrate lipid, the P loop needs to extract it from the bilayer, since the loop is located highly distal to the bilayer surface. Our data illustrate that PIP_3_ significantly protrudes from the bilayer surface and repositions toward the active site (Figure 5A). The protruding PIP_3_ can form salt bridges with key basic residues in the P loop. Deviations of the lipids from the bilayer surface show prominent protrusions of the lipids in the *holo* membrane side containing the protein as compared to those in the *apo* membrane side (Figure 5B). Note that PIP_2_ in *holo* also shows a weak protrusion but less significantly than PIP_3_, since PIP_2_ pursues the basic residues of PBD that reside in the bilayer surface. No significant protrusions of the phosphoinositide lipids can be observed for the monotonic systems containing only PIP_2_ (P2 system) or PIP_3_ (P3 system) (Figure S6A). This indicates that the P2 system is deficient in PIP_3_, and the P3 system fails to recruit PIP_3_ to the active site. In the PTEN phosphatase domain, the P loop contains the catalytic signature motif 123HCKAGKGR130 that is characteristic of the active site of the phosphatase family (Lee et al., 1999). It is important for Arg130 to coordinate PIP_3_ in the catalytic activity, since Arg130 facilitates migration of a phosphate group from the inositol head of PIP_3_ to the sidechain of Cys124. In addition to the catalytically significant Arg130, we discovered that two Lys residues, Lys125 and Lys128 in the P loop also coordinate PIP_3_. These Lys residues are PTEN-specific. In the P loop, two PIP_3_ are observed in the coordination; one with Lys128 and the other with Lys125 and Arg130. The PIP_3_ coordination changes the P loop to an “open” conformation with the rearrangement of the sidechains of the catalytic residues, Cys124 and Arg130, toward the substrate (Figure 5C). In the crystal structure of PTEN, the P loop shows a “collapsed” conformation (Figure S6B). In our simulations, the P loop retains the collapsed conformation in the PS, P2, and P3 systems (Figure S6C). The PS and P2 systems lack PIP_3_ in the bilayers, but the P3 system retains it. We found that the P3 system fails to coordinate PIP_3_ to Arg130 in the active site. The failure of the coordination of PIP_3_ to Arg130 is also observed in one PP system where the P loop retains the collapsed conformation. In PP systems with the PIP_3_ coordination, the sidechain of Asp92 in the WPD loop (residues 88-98) is rotated to the binding site, appearing to participate in the catalytic activity (Figure 5C and Figure S6D). This further implicates the “closed” WPD loop conformation with high catalytic activity (Brandao et al., 2012; Chia et al., 2010). However, studies of Asp92 mutations *in vivo* and *in vitro* suggested that Asp92 residue is not a catalytic amino acid commonly involved in the hydrolysis of PIP_3_ by PTEN (Rodriguez-Escudero et al., 2011).

**Figure 5.**
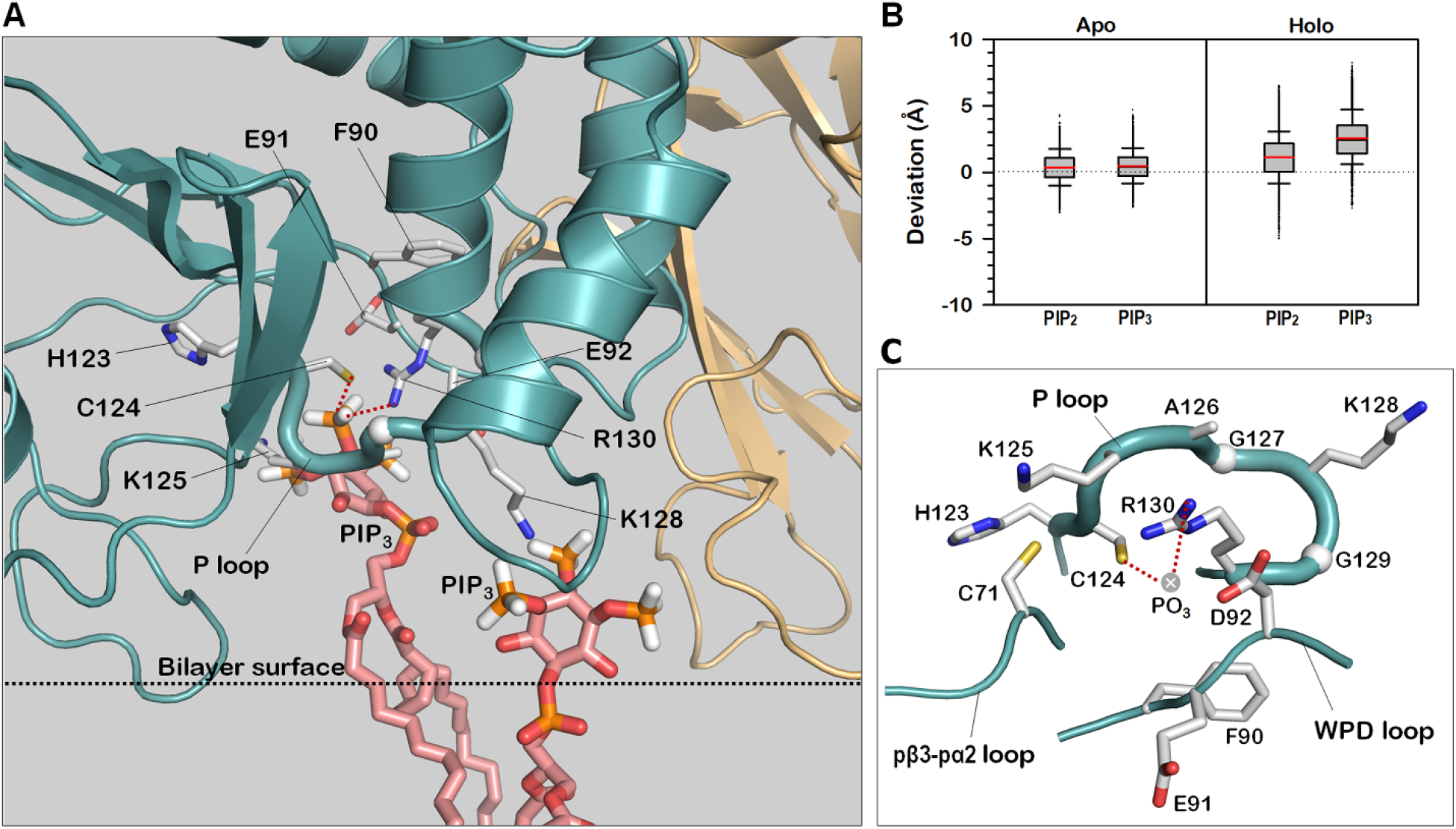
PIP_3_ coordination induces the P loop conformational change. (A) A snapshot highlighting P loop and protruded PIP_3_ from the bilayer surface at the active site. (B) Deviations of the phosphate atoms of PIP_2_ and PIP_3_ from the averaged position of the phosphate atoms of DOPC and DOPS for the *apo* and *holo* bilayer leaflets in the absence and presence of the protein, respectively. (C) The open conformation of P loop with the coordination of PIP_3_ at the active site.

In PTEN, the phosphatase and C2 domains are hinged by a short linker, 186LDYR189, located far from the bilayer surface. Underneath the linker, there is a tunnel-like, water-filled space encompassed by these two domains and the bilayer surface (Figure 6A). Hereafter, we refer the water-filled tunnel to aqueous canal. Through this region, aqueous molecules can penetrate the active site. Also, PIP_3_ with large inositol head can enter the region from both ends and move to the active site along the path connecting the PIP_3_-favored distal basic residues. The HOLE program (Smart et al., 1993) calculated the structure of the aqueous canal, illustrating that the canal’s dimensions are large enough for water transport (Figure 6B). Catalytic activity requires the water molecules for hydrolysis to release the phosphate group from Cys124 after transferring it from PIP_3_. Three-dimensional water density map illustrates high populations of residual waters around the sidechains of Cys124 and Arg130 at the active site (Figure 6C). The high probability of water suggests that the lipid substrate transports water molecules that soak the active site. In contrast, density maps with less waters populating the sidechains of catalytic residues in the P loop can be observed in the active sites of systems where PIP_3_ is absent (Figure S7). Due to formation of the water-filled space, no significant interdomain interaction between the phosphatase and C2 domains is observed (Figure 6D). However, the high residue-residue contact probability indicates that phosphatase residues Ser170 and Arg173 tightly hold the phosphatase–C2 domain interface underneath the short linker. Oncogenic mutations abolishing this interface (Smith et al., 2019) commonly occur at the interface residues Ser170 and Arg173, resulting in inactivation of PTEN in cancer (Lee et al., 1999). The N-terminal PBD marginally contacts the phosphatase domain, while the CTT largely contacts the phosphatase and C2 domains.

**Figure 6.**
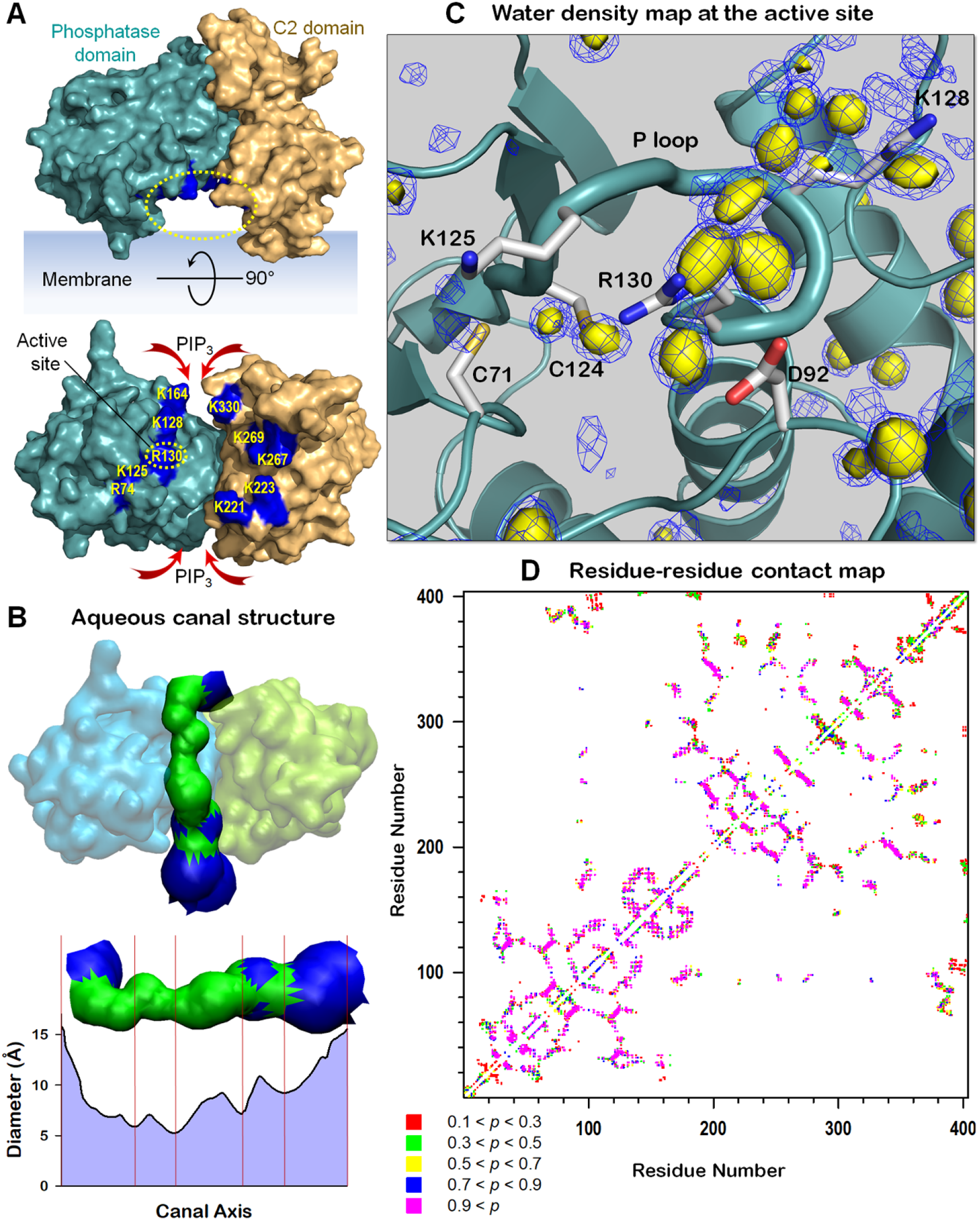
High density of waters at the active site for catalysis. (A) Highlight showing the tunnel-like, water-filled space formed by the inner surfaces of phosphatase and C2 domains and the bilayer surface denoting as the aqueous canal. Along the water-filled interface between the phosphatase and C2 domains, PIP_3_-favored distal basic residues are marked. (B) The structure and diameter of aqueous canal calculated with HOLE program (Smart et al., 1993). For the canal structure, green denotes the diameter in the range, 4 ≤ *d* ≤ 10 Å, and blue denotes the diameter of *d* > 10 Å. (C) Three-dimensional water density map with probabilities, *p* = 0.5 (yellow surface) and *p* = 0.4 (blue mesh). (D) Interdomain residue-residue contacts for PTEN. For two intramolecular residues *i* and *j*, the probability of contact for the distance between the 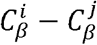 (C_α_ is used for Gly residue) with cutoff 10 Å was calculated. Residue *j* with |*j* – *i*| < 4 was omitted, and *p* < 0.1 was neglected in the calculation.

### Conformations of the IDR in the C2 domain and CTT in the C-terminal region

The IDR was initially unstructured, and most of the region in the domain remained disordered as it was. However, the segment 306IER308 participates in the β-sheet formation with cβ7 strand (Figure S8A). The β-sheet extension is stable as observed for most PTEN systems. Occasionally, the segment 299EIDSIC304 folds into an α-helix as calculated by STRIDE (Frishman and Argos, 1995) (Figure S8B), but most PTEN systems display it as a turn conformation. In contrast, the CTT retains the unstructured characteristics with large fluctuations in most PTEN systems. Its conformational space spans from a collapsed chain with radius gyration *R*_g_ = ∼15.0 Å to an extended chain with *R*_g_ = ∼24.0 Å (Figure S8C). The long unstructured CTT chain can reach the phosphatase domain as shown in the residue-residue contact map (Figure 6C), while the collapsed CTT mainly rides on the C2 domain. Thus, the interactions of CTT with the phosphatase and C2 domains are highly dynamic due to the various conformational ensembles. As observed in previous computational studies (Shenoy et al., 2012), no membrane interaction of CTT was observed, suggesting that CTT does not interfere in PTEN membrane localization and PIP_3_ recruitment by the phosphatase domain.

## Discussion

We report here selective membrane interaction of PTEN in the absence and presence of the signaling lipids phosphoinositide PIP_2_ and PIP_3_. The canonical role of PTEN is to negatively regulate the PI3K/AKT/mTOR pathway by dephosphorylating the lipid substrate PIP_3_ into PIP_2_. PTEN targets distinct membrane microdomains that are enriched in phosphoinositide lipids. Our studies demonstrate that effective membrane localization of PTEN depends on the lipid composition in the membranes. We mimic several types of microdomains within the plasma membrane (Table S1). In the zwitterionic membrane, PTEN abrogates its catalytic activity impairing the interaction of the phosphatase domain with the membrane, while the C2-membrane interaction remains intact. In the presence of phosphatidylserine, PTEN proceeds with membrane localization through the docking of the pβ2-pα1 loop of the phosphatase domain into the anionic membrane. The C2 domain conserves its intrinsic feature – targeting the cell membrane through the insertion of the CBR3 loop into both zwitterionic and anionic membranes. A series of basic Lys residues in the CBR3 loop buttress the interaction of PTEN with the anionic membrane containing phosphatidylserine. The process of PTEN membrane localization is completed through the stable anchorage of the unstructured PBD in the anionic membrane containing both phosphoinositides, PIP_2_ and PIP_3_ (Figure 7), suggesting that full activation of PTEN requires both lipids interacting with the specific domains; PIP_3_ at the active site and PIP_2_ at the PBD binding site in good agreement with experimental data (Campbell et al., 2003). We suggest that the coordination of PIP_3_ to the P loop in the active site of the phosphatase domain can allosterically promote PBD unfolding. Once the unfolded PBD recognizes PIP_2_ at the binding site, it is released from the phosphatase domain and translocated onto the membrane surface. Taken together, PTEN uses three major anchor points, the N-terminal PBD, the pβ2-pα1 loop in the phosphatase domain, and the CBR3 loop in the C2 domain to maintain the stable membrane-interacting conformation.

**Figure 7.**
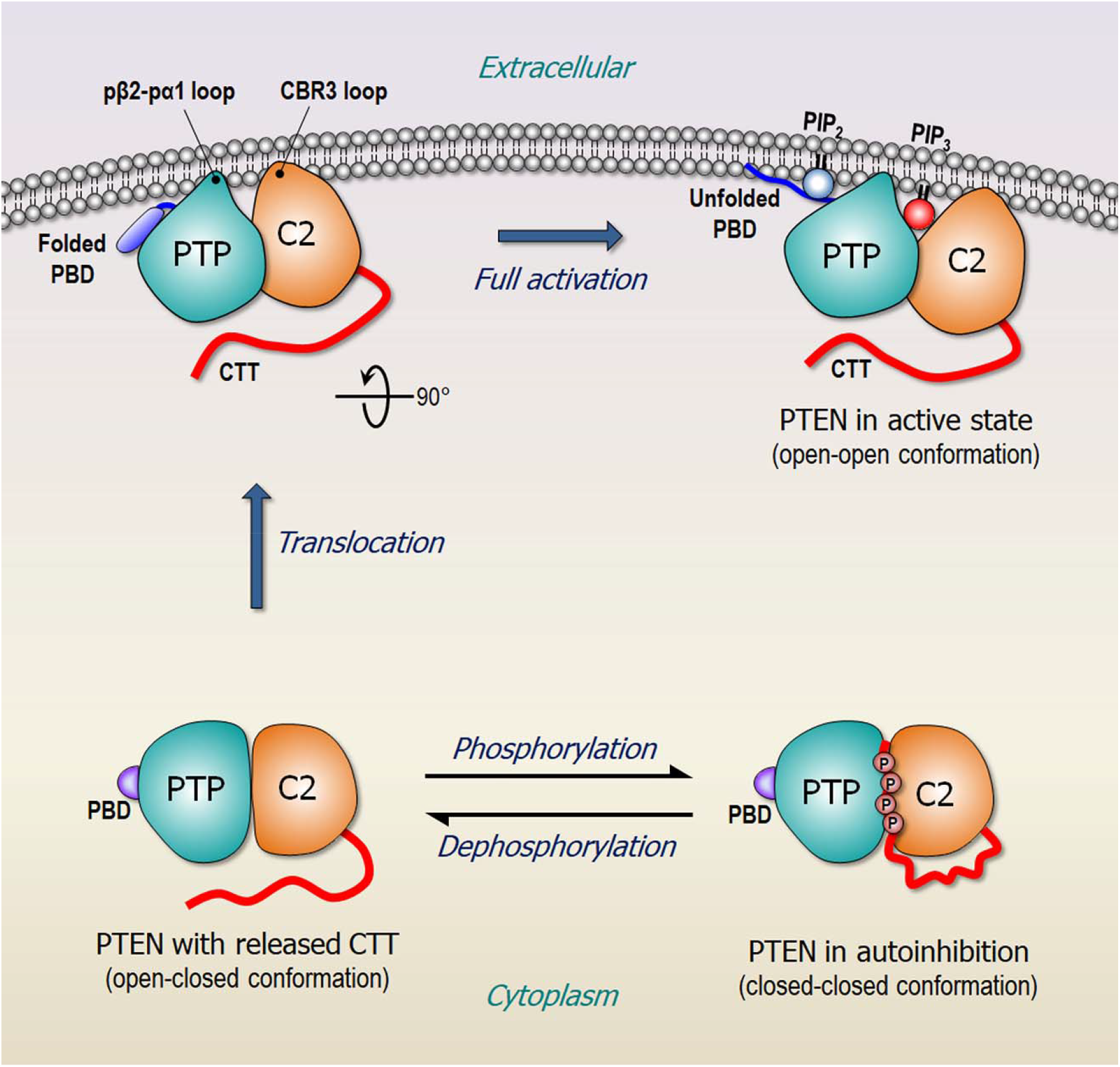
A schematic diagram illustrating membrane localization of PTEN. In the cytosol, PTEN is autoinhibited by the phosphorylated CTT, through the “closed-closed” conformation. Dephosphorylation on the CTT removes the autoinhibition, shifting the populations toward the “open-closed” conformation. With the exposed membrane-binding surface, PTEN can localize at the anionic membrane. Full activation of PTEN requires the membrane anchorage of PBD at the membrane microdomain enriched by both phosphoinositides, PIP_2_ and PIP_3_. In the cartoon, PTP denotes the phosphatase domain.

Substantial biophysical studies of PTEN membrane interactions pointed a significant role of PIP_2_ in the PTEN activity. As a product of PTEN, PIP_2_ interacting with PBD facilitates PTEN hydrolysis of PIP_3_, creating a positive feedback loop (Campbell et al., 2003). Our simulations also verified the requirement of both phosphoinositides in PTEN activation. It was reported that among phosphoinositides, PIP_2_ exhibits specific binding preference for both full-length PTEN and the N-terminal peptide PTEN_1-21_, but not for the N-terminal truncated PTEN_16-403_ (Redfern et al., 2008). This indicates that PTEN’s N-terminal PBD preferentially binds PIP_2_, consistent with our finding. In the simulations, our PTEN systems contain two types of phosphoinositides, PIP_2_ and PIP_3_, and we only observed the PBD interacting with PIP_2_, but not with PIP_3_. K13E mutation in PBD abolishes the ability of PTEN to bind anionic lipid bilayer (Walker et al., 2004). Recent ^31^P field cycling NMR spectroscopy on spin-labeled protein identified Lys13 and Arg47 as the PIP_2_ binding sites, which are distant from the active site (Wei et al., 2015). They discovered that PIP_2_ interactions at the binding sites secure the electrostatic anchorages of PTEN at the membrane facilitating PTEN processive catalysis. For the basic residues at the binding sites, our simulations showed high probability of forming salt bridges with PIP_2_, verifying the PIP_2_ binding sites. The PBD contains the polybasic patch with three arginine, Arg11, Arg14, and Arg15, and one lysine, Lys13. These Arg residues in the unfolded PBD exhibited high probability of forming salt bridges with PIP_2_. Similarly, Arg47 and Arg161 in the loops of phosphatase domain, and Arg335 in the loop of C2 domain also showed high probability of forming salt bridges with PIP_2_, suggesting that PIP_2_ preferentially binds Arg residues at the unstructured PTEN regions located nearby the membrane surface.

Previous computational studies indicated that electrostatic interactions initially drive PTEN to the membrane, followed by reorientation of the protein to optimize its interactions with the substrate (Kalli et al., 2014; Lumb and Sansom, 2013; Shenoy et al., 2012). The MD simulations combined with the neutron reflection (NR) experiments revealed that Arg161 and Lys163 in the phosphatase domain, and Lys330 and Lys332 in the C2 domain have considerable lipid residence time for phosphoinositide lipids, but not for DOPC and DOPS lipids (Nanda et al., 2015; Shenoy et al., 2012). In our simulations, these residues exhibited high probability of forming salt bridge with the phosphoinositides, PIP_2_ and PIP_3_. The profile of peaks in the lipid residue time (Nanda et al., 2015) is similar to that in the lipid contact probability from our simulations, suggesting that PTEN aligns at the membrane in a similar way. They further noticed that the phosphatase domain diffuses into the membrane slightly deeper in the presence of phosphoinositides than that in the phosphoinositide-free membrane. Our simulations also demonstrated that PTEN systems containing phosphoinositides (P2, P3, and PP systems) have smaller deviations of phosphatase domain from the bilayer surface than the PC and PS systems without phosphoinositides. In the PS system, individual PTEN domains from the bilayer surface are similarly located as observed in the PTEN system with the DOPC:DOPS bilayer in 2:1 molar ratio (Shenoy et al., 2012), especially for the elevated PBD location. However, in the presence of both PIP_2_ and PIP_3_, the PBD spontaneously translocated onto the membrane surface targeting PIP_2_. The CTT exhibits a random conformation, thoroughly avoiding membrane interaction during the simulations in agreement with previous computations studies (Shenoy et al., 2012). Our data indicate that the salt bridge interactions between the basic residues and the lipids are a major driving force in stabilizing the protein-membrane interface. These basic residues are in the three major anchor points among the domains, located close to the membrane surface with the proximal basic residues forming salt bridges with the phosphate atom of the head groups of phosphatidylcholine and phosphatidylserine. In contrast, the basic residues distal to the membrane are positioned at the water-filled interface between the phosphatase and C2 domains. We denoted the tunnel-like, water-filled space formed by the inner surfaces of these domains and the membrane surface as the aqueous canal. The distal basic residues including the catalytically significant Lys125, Lys128, and Arg130 in the P loop, are closely associated with PIP_3_ migration towards the active site. Lys125 occasionally coordinates PIP_3_ together with Arg130, suggesting its auxiliary role in the catalysis, and Lys128 has high probability of PIP_3_ binding. Other distal basic residues in the aqueous canal may pave the road for the substrate lipid to move toward the active site, serving as a reservoir for the PIP_3_-binding sites. Furthermore, the open P loop conformation points to PTEN in the active state, primed for catalysis. Importantly, mutations and acetylation of Lys125 and Lys128 result in a reduction of PTEN phosphatase activity (Lee et al., 1999; Okumura et al., 2006) further demonstrating their critical role in catalysis. Overall, observations of PIP_3_ coordination at Cys124 and Arg130, the proximity of Asp92 in the WPD loop to the coordination, and the high population of the residual waters at the active site suggest that these events participate in cooperative catalysis.

Previous PTEN studies showed that affinity to the membrane is reduced by the following: cancer-associated mutations (Masson and Williams, 2020) inhibit recruitment to the plasma membrane, thus abolishing its interactions with phospholipids (Nguyen et al., 2015); phosphorylation of the CTT serine-threonine cluster impairs PTEN membrane binding ability, resulting in a closed protein conformation (Bolduc et al., 2013; Malaney et al., 2013; Masson and Williams, 2020; Rahdar et al., 2009; Ross and Gericke, 2009); CTT deletion decreases protein stability leading to tumor development (Georgescu et al., 1999; Sun et al., 2014); and a mutation in the PBD (K13E) leads to PTEN inactivation (Masson and Williams, 2020; Nguyen et al., 2015; Walker et al., 2004). In the cytosol, monomeric PTEN is in the inactive state with its PBD bound to the phosphatase domain and the phosphorylated CTT interfering with the membrane binding interface (Bolduc et al., 2013; Chen et al., 2016; Malaney et al., 2013; Masson et al., 2016; Masson and Williams, 2020; Mingo et al., 2019; Rahdar et al., 2009; Ross and Gericke, 2009; Vazquez et al., 2000) (Figure 7), whereby the inactive, autoinhibited PTEN is in the “closed-closed” conformation.

In the autoinhibited PTEN conformation proposed earlier, the long unstructured phosphorylated CTT mediates the suppression of PTEN catalytic activity, covering the C2 domain membrane binding interface (Malaney et al., 2013; Odriozola et al., 2007; Rahdar et al., 2009; Ross and Gericke, 2009). This prevents the distal basic residues positioned along the aqueous canal recruiting the substrate. We suspect that the highly negatively charged IDR at the opposite side of C2 promotes the ‘correct’ interaction of the phosphorylated CTT with the membrane binding interface, spawning the autoinhibited state. Dephosphorylation of the CTT shifts the population to the membrane binding-ready, “open-closed” conformation. With the exposure of its membrane binding interface, PTEN can localize on the membrane. In the absence of phosphoinositides, the PBD is still in a closed conformation. However, PTEN acquires full activation with the “open-open” conformation, when interacting with membrane microdomains enriched with phosphoinositides. Notably, PTEN also targets the membrane microdomains that PI3K does, co-localizing with its substrate, suggesting that both negative and positive regulators in the PI3K/AKT/mTOR pathway concomitantly exist at the same microdomains within the plasma membrane.

To date, PTEN has been considered a major clinically undruggable target due to its status as a tumor suppressor phosphatase and its complex regulation. A major challenge in thwarting its oncogenic mutations is that PTEN is a tumor suppressor, not an activator. That is, oncogenic PTEN is the inactive; not the active form. Activation is dramatically more difficult to achieve than inactivation since the active site residues must be precisely positioned for catalysis and the regulatory machinery appropriately tuned. With the diverse mutations these vary in the inactive state. Nevertheless, the mechanistic details elucidated in this work may suggest exploring at least two potential venues. The first relates to relieving PTEN’s autoinhibition, shifting from an inactive state to an active state. Though this has been previously explored (Kalli et al., 2014; Lumb and Sansom, 2013), the details provided here may suggest additional PTEN structural features to target. We suggest that charge reversal mutations on the IDR release the autoinhibition through the interaction with the phosphorylated CTT, exposing the membrane binding interface of the phosphatase and C2 domains and thus promoting membrane localization of PTEN. The second relates to the design of allosteric drugs that restore the coordination in PTEN mutants in which catalysis is impaired, such as the C124S mutant which is catalytically dead (Bonneau and Longy, 2000), and the G129E mutation which inhibits the recognition of phosphoinositide (Myers et al., 1998). Here we detail the substrate coordination and conformational change of the P loop that implicate PTEN catalytic activity. We observed that PIP_3_ significantly protrudes from the membrane surface and can form salt bridges with key basic residues in the P loop. PIP_2_ in the *holo* state also protrudes but less significantly than PIP_3_ does. Residues Cys124 and R130 are essential for catalysis whereby is vital for PIP_3_ coordination, facilitating the phosphate group migration from its inositol to Cys124. Lys125 and Lys128 in the P loop also coordinate PIP_3_ and are PTEN-specific. Moreover, we observed two crucial active site coordination events, namely PIP_3_ with Lys128, and Lys125 and Arg130, which alters P loop conformation through sidechain rearrangement of the catalytic residues, Cys124 and Arg130, toward the substrate. The use of allosteric modulators to restore these coordination events in PTEN mutants can provide therapeutic advantages. One possible approach would be via the design of an allostery-based rescue mutant as we have proposed for PI3Kα (Zhang et al., 2020) and indeed PTEN (Nussinov et al., 2020), and earlier for the von Hippel-Lindau tumor suppressor protein (Liu and Nussinov, 2008). An alternative approach to overcome any mutational effects and restore wild-type PTEN function would be the design of small-molecule allosteric modulators that target the allosteric communication pathway from the PBD region to the active site that stabilizes PTEN–membrane interactions.

In conclusion, membrane localization and orientation of PTEN strongly depend on the lipid composition of the membrane. Our studies elucidate the mechanistic details of PTEN activation at the membrane and the role of phosphoinositides in the conformational changes of membrane bound PTEN. Cancer-associated mutations prevent effective membrane localization, leading to the dysfunctional inhibition of oncogenic cell signaling and growth. With PTEN being one of the most frequently mutated proteins in cancers, obtaining its membrane associated structure and mechanism in atomic resolution may provide insight assisting drug discovery, for the currently still undruggable major cancer target (McLoughlin et al., 2018).

## Acknowledgments

This study was funded, in part, by the Ambrose Monell Foundation (to C.E.). This project has been funded in whole or in part with federal funds from the National Cancer Institute, National Institutes of Health, under contract HHSN26120080001E. The content of this publication does not necessarily reflect the views or policies of the Department of Health and Human Services, nor does mention of trade names, commercial products, or organizations imply endorsement by the U.S. Government. This Research was supported [in part] by the Intramural Research Program of the NIH, National Cancer Institute, Center for Cancer Research. All simulations had been performed using the high-performance computational facilities of the Biowulf PC/Linux cluster at the National Institutes of Health, Bethesda, MD (https://hpc.nih.gov/).

## Author contributions

H.J. and R.N. conceived and designed the study. I.N.S. prepared the initial protein model, and H.J. conducted most of the simulations, analyzed the results, and wrote most of the paper. H.J., I.N.S., C.E., and R.N. have given approval to the final version of the manuscript.

## Declaration of interests

The authors declare no competing interests.

## Method details

### Preparing the full-length PTEN

The crystal structure of wild-type PTEN (PDB: 1D5R) was used to model the full-length PTEN protein. The missing coordinates for residues 1-13 in PBD, residues 282-312 of IDR in C2, and residues 352-403 in the CTT were obtained by a hierarchal iterative template-based threading approach using the I-TASSER program (Roy et al., 2010; Yang et al., 2015; Yang and Zhang, 2015). The PBD was modelled as a small α-helix, and the IDR and CTT were modelled as unstructured chains (Figure 1). *In silico* modelling of full-length PTEN showed that the α-helix PBD interacts with the phosphatase domain and both unstructured IDR and CTT chains span the phosphatase and C2 domains marginally interacting with them. We ensured that the long disordered CTT chain does not block the catalytic site at the membrane binding interface. In autoinhibition, the catalytic site is blocked by phosphorylated serine-threonine cluster (Ser380, Thr382, Thr383, and Ser385) in the CTT, which is crucial for maintaining PTEN in a stable cytosolic state (Masson and Williams, 2020).

### Construction of the bilayer systems with PTEN

To generate the initial configuration of membrane bound PTEN, the protein was placed on the lipid bilayer without contacting the lipids. In the initial placement, PTEN faced its putative membrane-binding interface toward the bilayer surface as suggested by the crystal structure (Lee et al., 1999). In this orientation, the pβ2-pα1 loop (residues 40-47) in the phosphatase domain and the CBR3 loop (residues 260-269) in the C2 domain were closely located at the bilayer surface. The initial construction revealed that all modelled regions including the N-terminal PBD do not interact with the bilayer before starting of the simulations. An equilibrated and hydrated lipid bilayer was constructed at the room temperature ensuring that the lipid bilayer is in the liquid phase. In the construction, preequilibrated lipids from the lipid library containing a couple of thousand lipids with different individual conformations generated from the CHARMM program (Brooks et al., 2009) were extracted and replaced with the pseudospheres interacting through the van der Waals (vdW) force field at the putative locations for the phosphate atom of lipid head group (Woolf and Roux, 1994, 1996). The bilayer contained 4 different lipids including the zwitterionic DOPC, anionic DOPS, and the phosphoinositide PIP_2_ and PIP_3_ lipids in 32:6:1:1, molar ratio (denoted ad PP system, Table S1). Total of 400 lipids constituted the bilayer system with lateral cell dimension of ∼120 Å^2^ for the unit cell. TIP3P waters were added at both sides of the bilayer with lipid/water ratio of ∼1/130.

### Atomistic molecular dynamics simulations

We performed MD simulations using the updated CHARMM (Brooks et al., 2009) all-atom force field (version 36m) (Huang et al., 2017; Klauda et al., 2010) for constructing the set of starting points and relaxing the systems to a production-ready stage. Our simulations closely followed the same protocol as in our previous works (Jang et al., 2015; Jang et al., 2017; Jang et al., 2016a; Jang et al., 2016b; Jang et al., 2020; Zhang et al., 2019a, b). The bilayer system containing PTEN and solvent has almost 210,000 atoms. For reproducibility, we constructed four independent systems with different lipid arrangements around protein. Statistics were taken over these trajectories. To observe the effects of lipid compositions on the PTEN-membrane interaction, four additional systems were constructed, including a pure DOPC bilayer (PC system), an anionic DOPC:DOPS (4:1, molar ratio) bilayer (PS system), and two anionic DOPC:DOPS:PIP_2_ (P2 system) and DOPC:DOPS:PIP_3_ (P3 system) bilayers in 16:3:1, molar ratio (Table S1). The former and latter phosphoinositide lipid bilayers radically contained PIP_2_ and PIP_3_, respectively.

In the preequilibrium stage, we performed a series of minimization and dynamics cycles for the solvents including lipids around the harmonically restrained protein. Each bilayer system was subjected to the preequilibrium simulation for 5 ns with the restrained backbones of PTEN until the solvent reached 310 K. The harmonic restraints on the backbones of PTEN were gradually removed through the solvent dynamics cycles with the particle mesh Ewald (PME) electrostatics calculation (Darden et al., 1993). The final dynamics cycle without the harmonic restraints completed the preequilibrium stage, generating the starting point for the production run. A total of 8.0 μs simulations were performed for the 8 systems, each with 1 μs. In the production runs, we employed the Langevin temperature control (Wu and Brooks, 2003) to maintain the constant temperature at 310 K and the Nosé-Hoover Langevin piston pressure control to sustain the pressure at 1 atm. To constrain the motion of bonds involving hydrogen atoms, we applied the SHAKE algorithm (Ryckaert et al., 1977). The production runs were implemented with the NAMD parallel-computing code (Phillips et al., 2005) on a Biowulf cluster at the National Institutes of Health (Bethesda, MD). In the analysis, the first 200 ns trajectories were removed, and thus averages were taken afterward.

## Notes

### Competing Interest Statement

The authors have declared no competing interest.

### Summary of Updates

Discussion section is revised.

