## Supplemental figures and a table for "The mechanism of full activation of tumor suppressor PTEN at the phosphoinositide-enriched membrane"

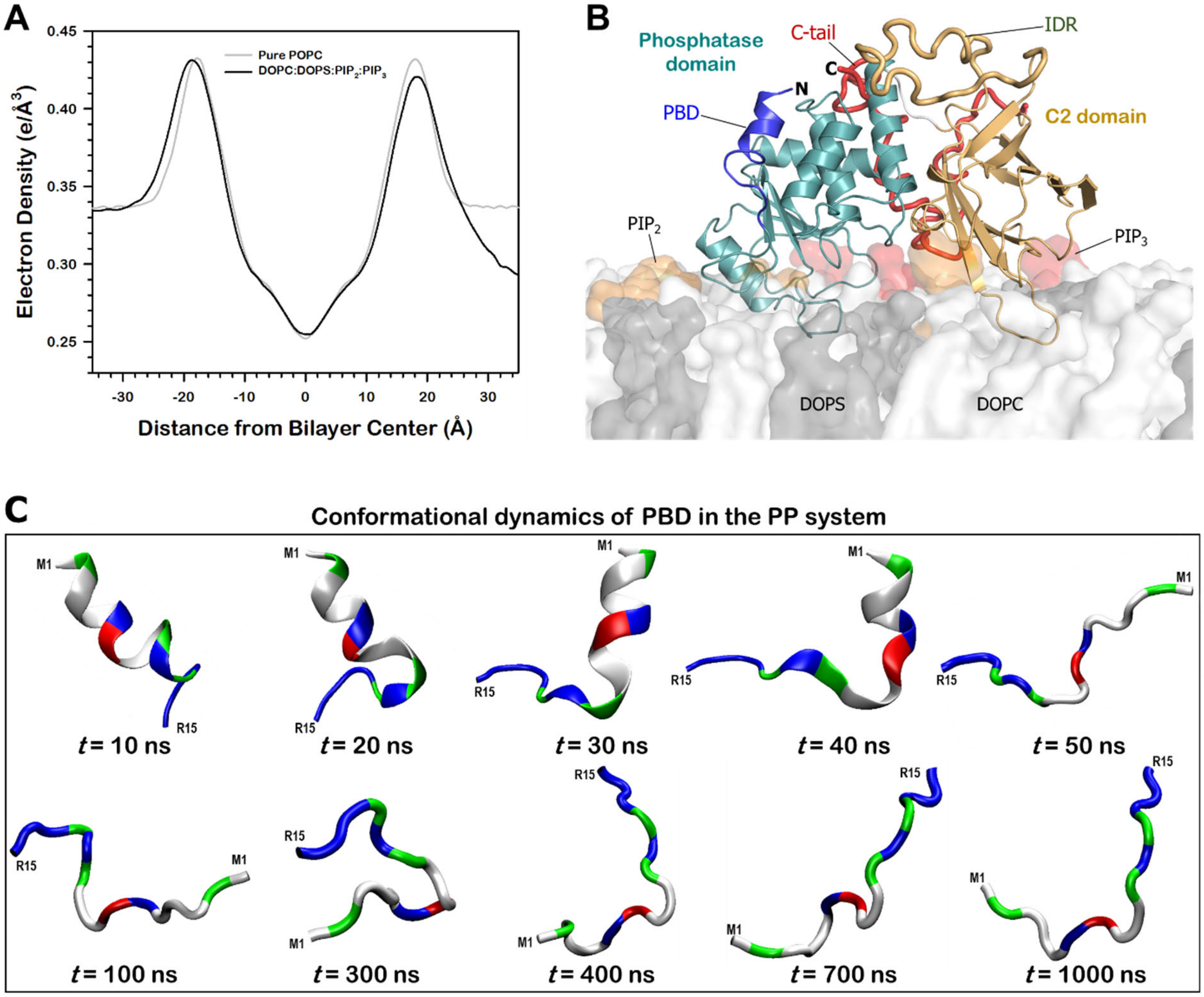


**Figure S1. Initial configuration of the PTEN-membrane system**, Related to Figure 2

(A) Electron density across the bilayers for the pure DOPC bilayer (gray line) and the PP system (black line) with the lipid composition of DOPC:DOPS:$PIP_2$:$PIP_3$ = 32:6:1:1, in molar ratio.

(B) A snapshot representing the initial conformation of PTEN in the PP system.

(C) Time-evolution structures of PBD in the PP system. In the cartoons, hydrophobic, polar/glycine, positively charged, and negatively charged residues are colored white, green, blue, and red, respectively.

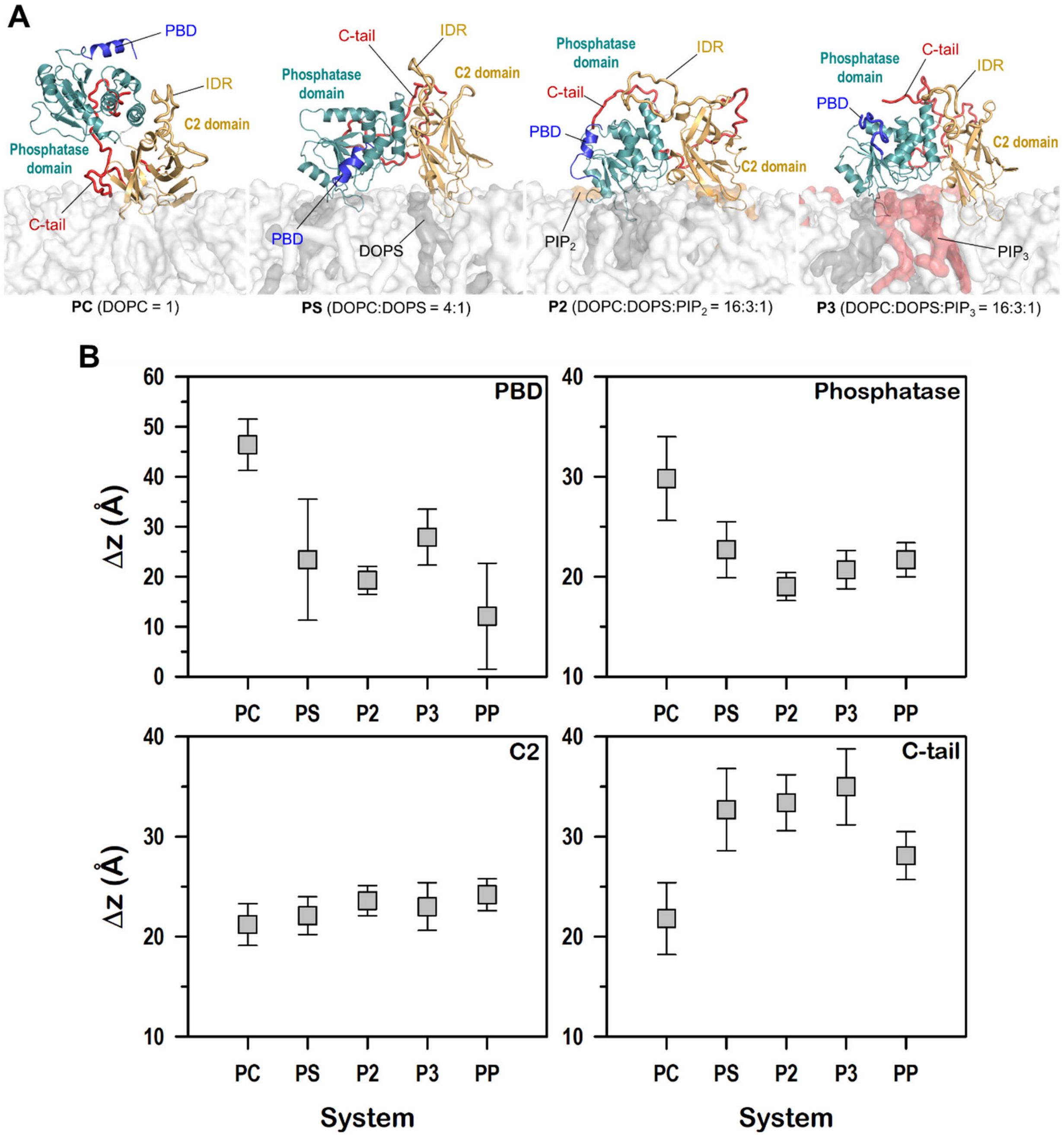


**Figure S2. Relaxed PTEN conformations in different lipid bilayer systems**, Related to Figure 3

(A) Snapshots representing the final conformations of PTEN in the PC, PS, P2, and P3 systems.

(B) Averaged deviations of the center of mass of individual PTEN domains from the bilayer surface for different bilayer systems.

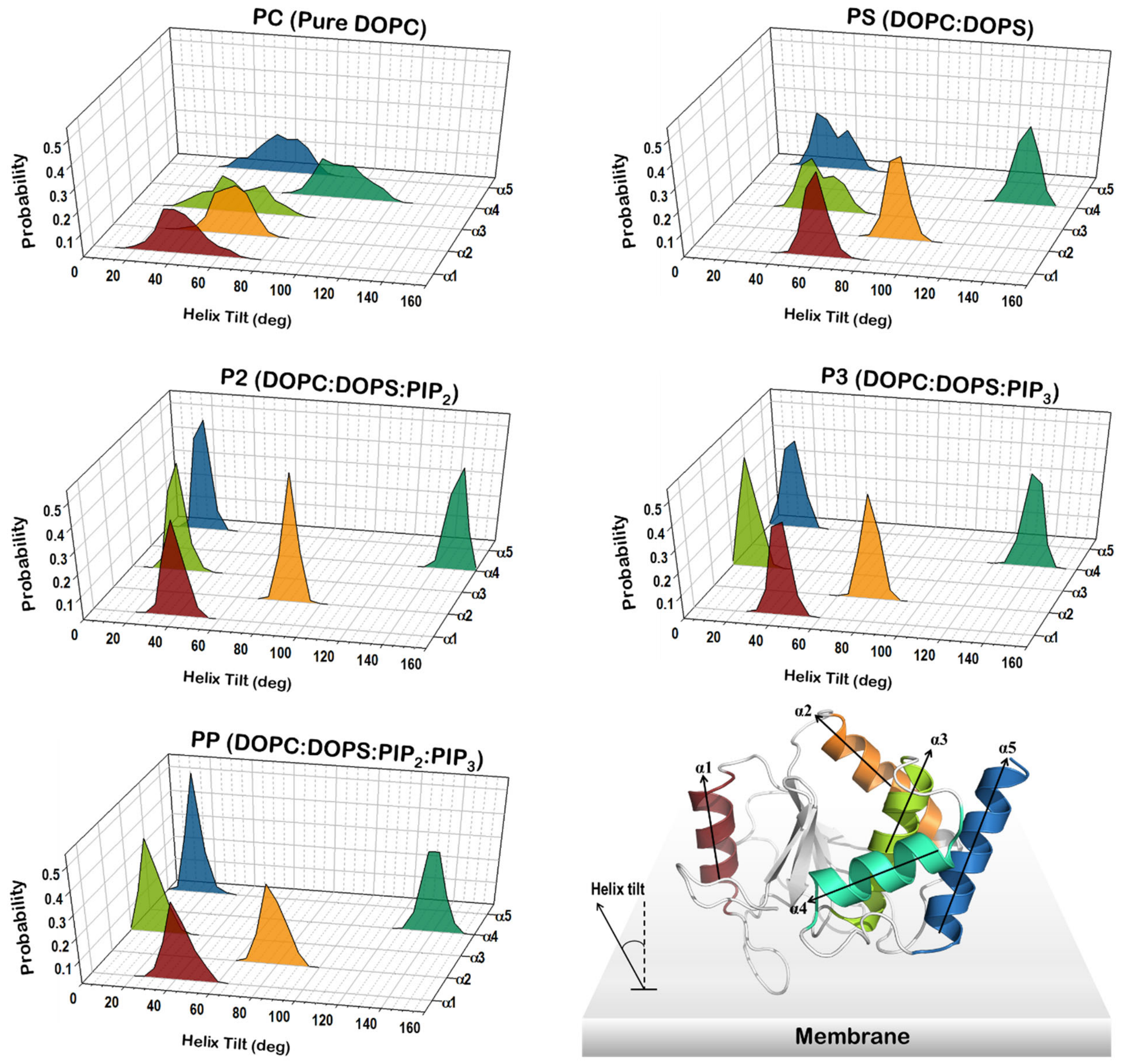


**Figure S3. Helix tilt angle of PTEN**, Related to Figure 3.
Probability distribution functions of the helix tilt with respect to the bilayer normal for helices in the phosphatase domain of PTEN in different bilayer systems.

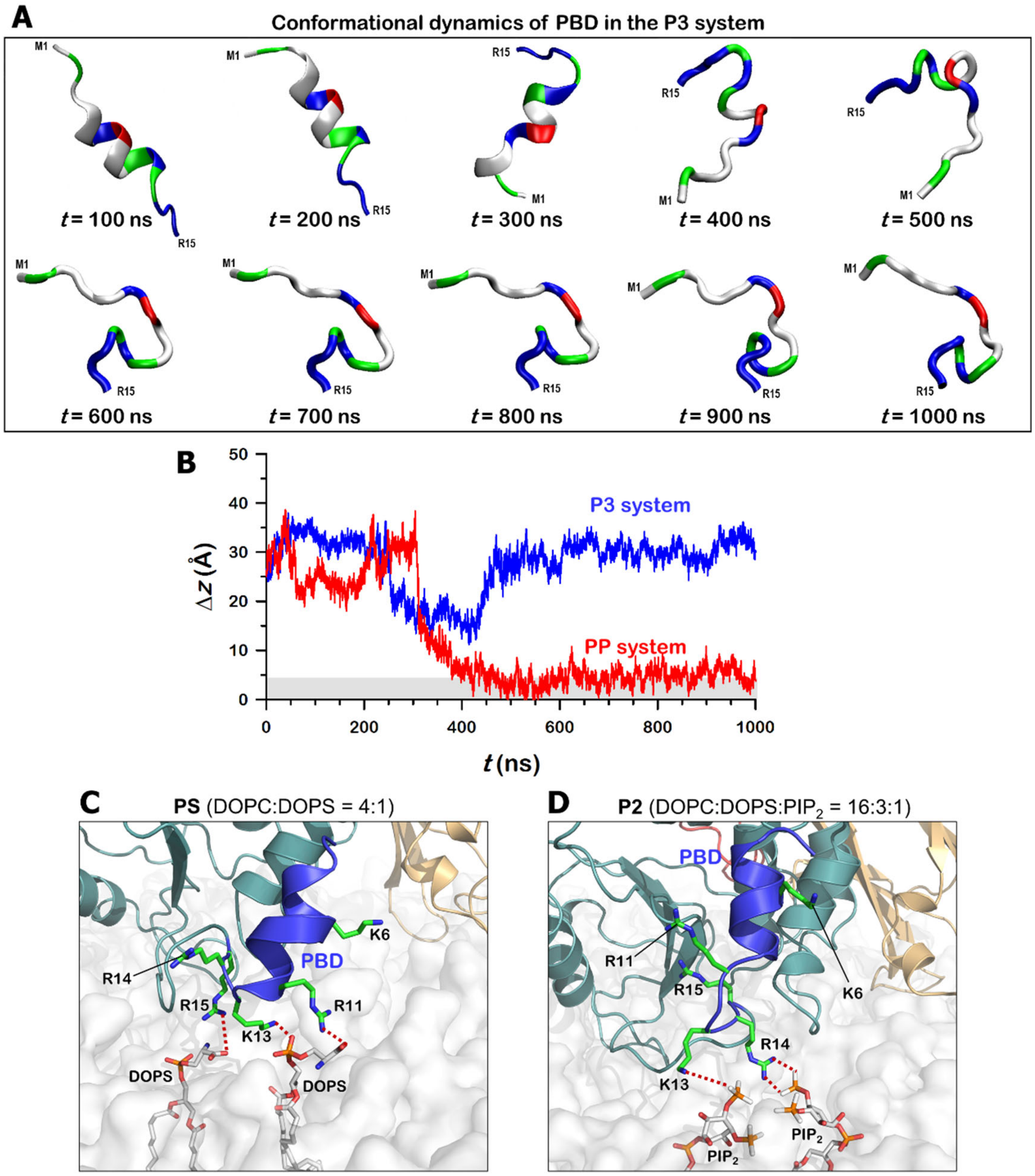


**Figure S4. Conformational dynamics of PBD in different bilayer systems**, Related to Figure 3

(A) Time-evolution structures of PBD in the P3 system. In the cartoons, hydrophobic, polar/glycine, positively charged, and negatively charged residues are colored white, green, blue, and red, respectively.

(B) Time series of the deviations of the center of mass of PBD in the P3 system compared to that in the PP system.

(C) Highlights shown the interactions of folded PBDs with the lipids in the PS system.

(D) The same for the P2 system. Key basic residues, Lys6, Arg11, Lys13, Arg14, and Arg15 are marked.

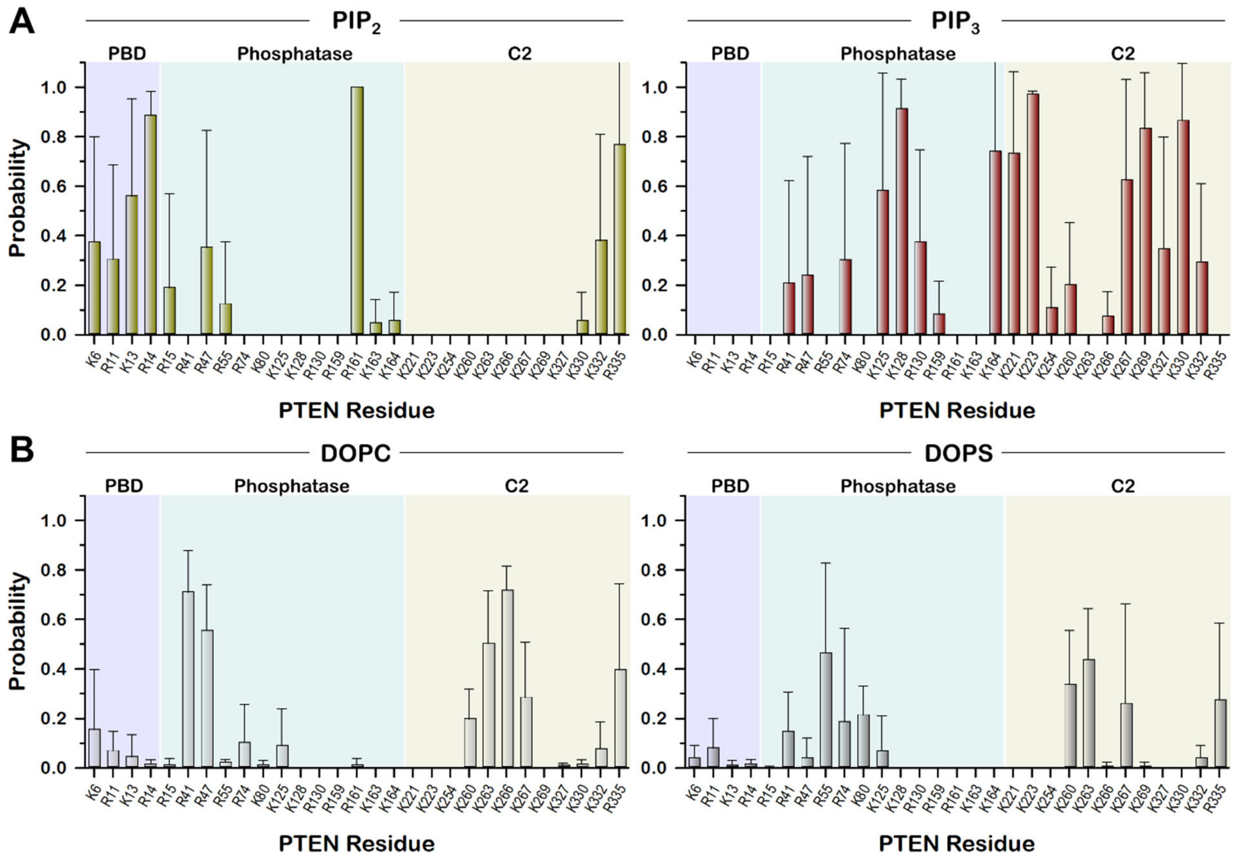


**Figure S5. Salt bridge interactions of key basic residues**, Related to Figure 4

(A) Two-dimensional bar graphs with error bars of the probabilities of salt bridge formation between the key basic residues at the membrane-binding interface and phosphoinositides, $PIP_2$ and $PIP_3$.

(B) The same for bulky lipids, DOPC and DOPS. Salt bridge is calculated between the nitrogen atoms in the sidechains of basic residues and the oxygen atoms in the phosphate group of all lipids and the inositol head group of phosphoinositides with the cutoff distance of 3.2 Å.

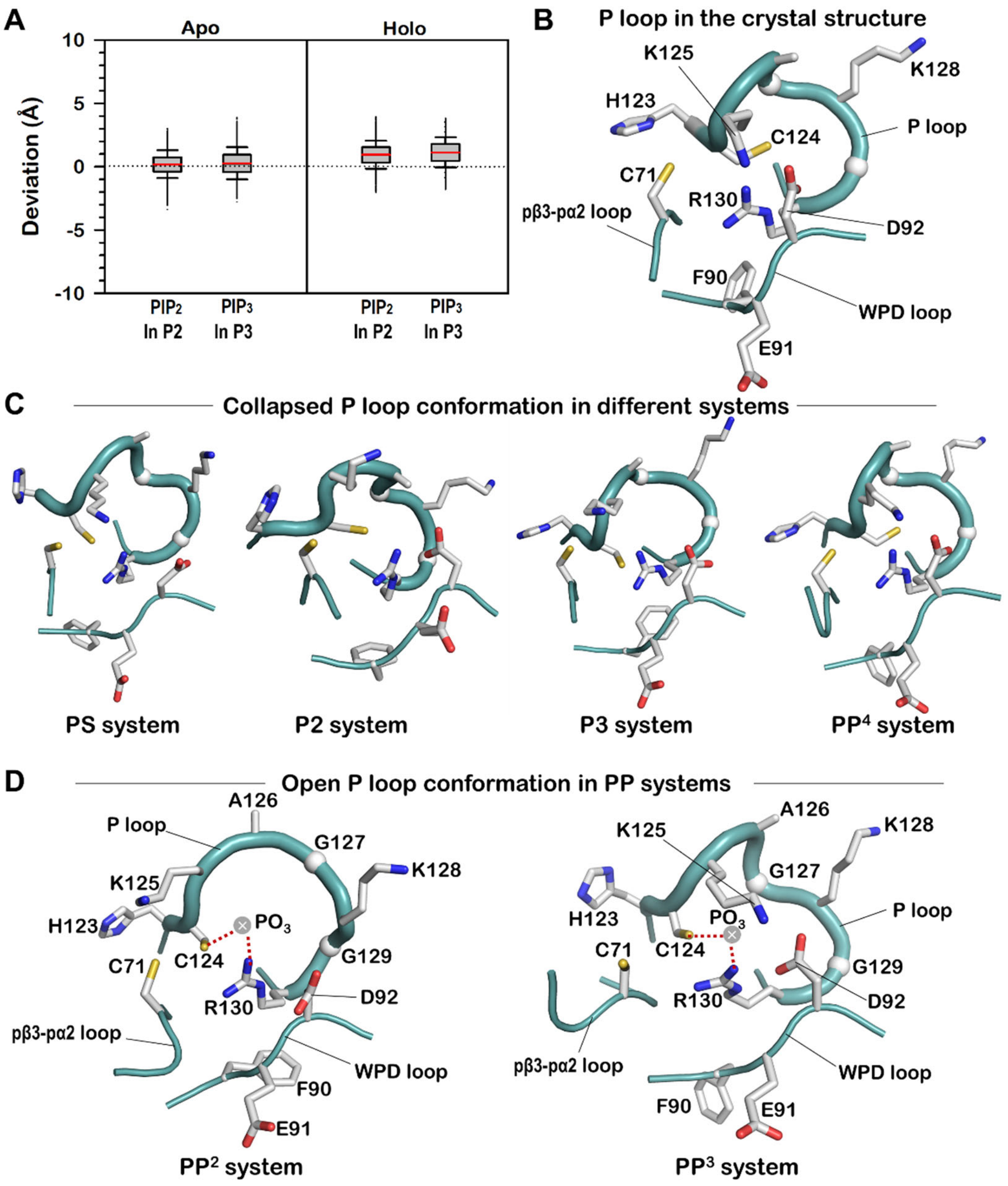


**Figure S6. Conformational change of P loop in different bilayer systems**, Related to Figure 5

(A) Deviations of the phosphate atoms of $PIP_2$ in the P2 system and $PIP_3$ in the P3 system from the averaged position of the phosphate atoms of DOPC and DOPS for the *apo* and *holo* bilayer leaflets in the absence and presence of the protein, respectively.

(B) A snapshot representing the P loop conformation in the crystal structure.

(C) Snapshots representing the collapsed P loop conformations observed in the PS, P2, P3, and $PP^4$ systems. One of the PP systems, $PP^4$ yields the collapsed P loop due to failure of the coordination of $PIP_3$ to Arg130.

(D) Snapshots representing the open conformation of P loop with the coordination of $PIP_3$ at the active site for the PP systems.

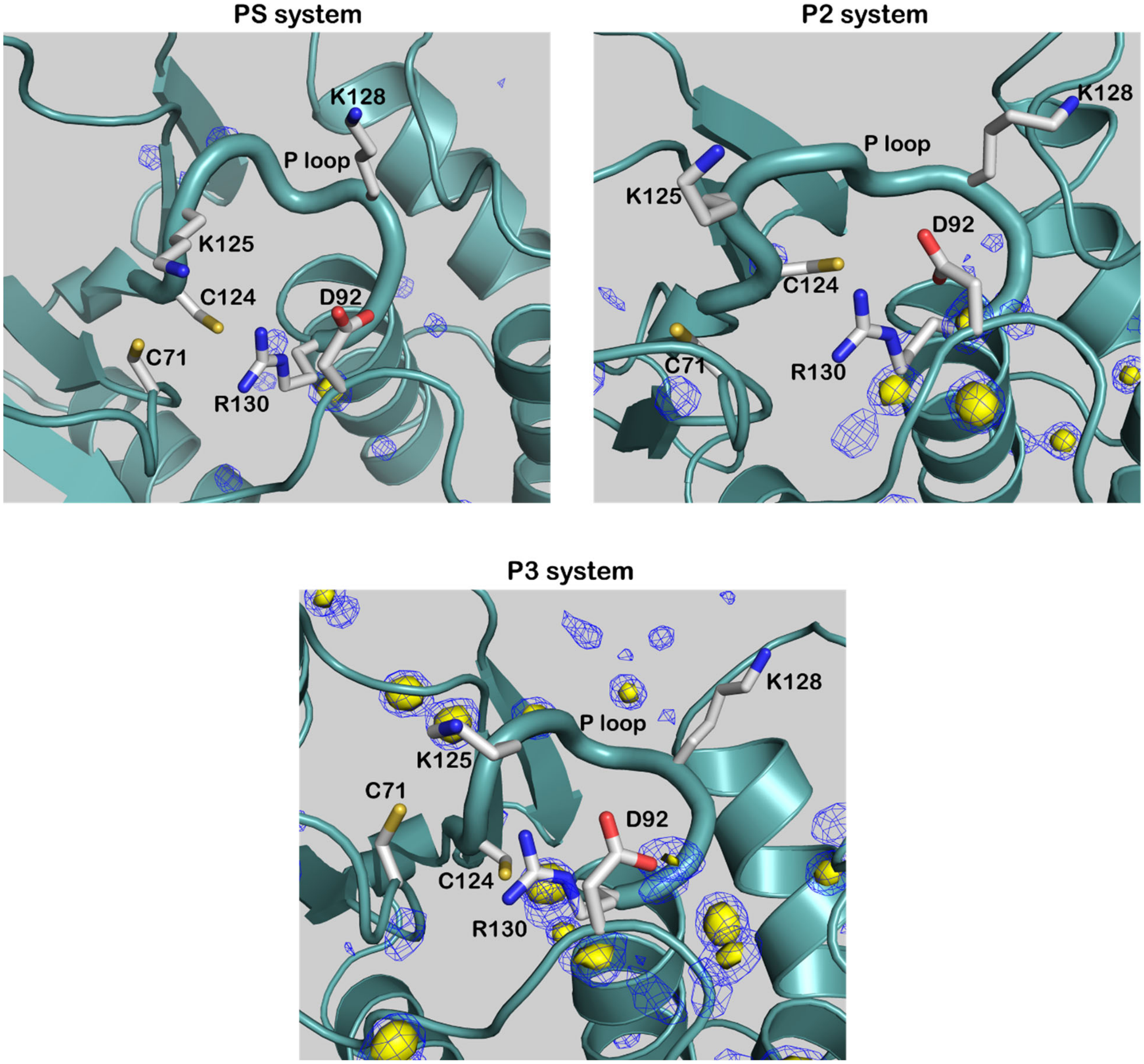


**Figure S7. Density of waters at the active site in different bilayer systems**, Related to Figure 6
Three-dimensional water density map with probabilities, $p = 0.5$ (yellow surface) and $p = 0.4$ (blue mesh) for the PS, P2, and P3 systems.

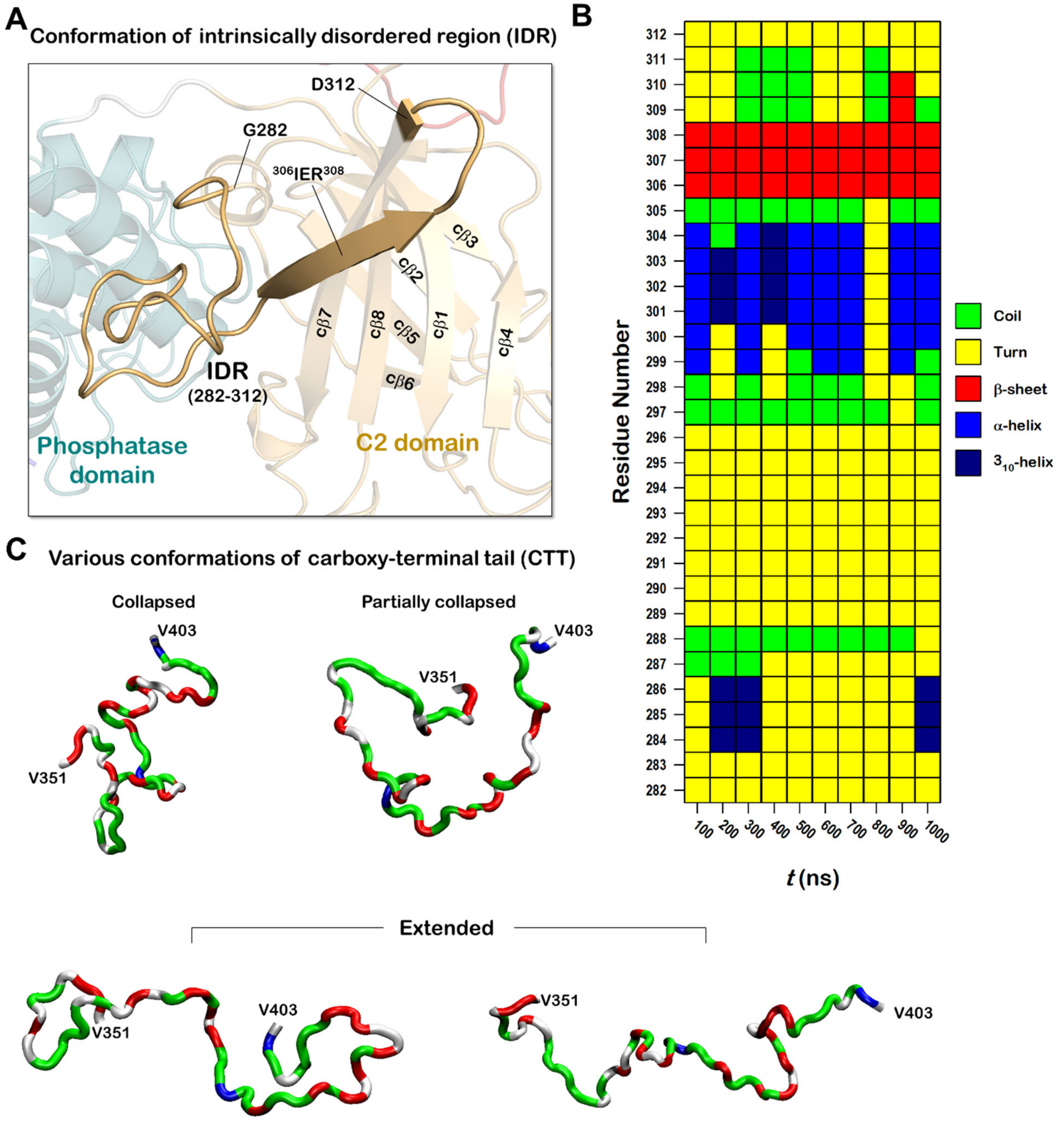


**Figure S8. Conformations of IDR and CTT**, Related to Figure 6

(A) Snapshot of the IDR conformation with the β-sheet extension.

(B) Descriptions of the secondary structure by the STRIDE program (Frishman and Argos, 1995) for the IDR depicted each 100 ns interval.

(C) Representative conformations of the CTT with the collapsed and extended chain structures.

**Table S1.** Composition of the lipid bilayer systems in the simulations. Related to STAR Methods

| System | Charged state | # of DOPC | # of DOPS | # of $PIP_2$ | # of $PIP_3$ | Molar ratio |
|---|---|---|---|---|---|---|
| **PC** | Zwitterionic | 400 | - | - | - | 1 |
| **PS** | Anionic | 320 | 80 | - | - | 4:1 |
| **P2** | Anionic | 320 | 60 | 20 | - | 16:3:1 |
| **P3** | Anionic | 320 | 60 | - | 20 | 16:3:1 |
| **PP** | Anionic | 320 | 60 | 10 | 10 | 32:6:1:1 |
